# Considering Zeros in Single Cell Sequencing Data Correlation Analysis

**DOI:** 10.1101/2023.05.13.540566

**Authors:** Guoshuai Cai, Xuanxuan Yu, Feifei Xiao

## Abstract

Single-cell sequencing technology has enabled correlation analysis of genomic features at the cellular level. However, high levels of noise and sparsity in single-cell sequencing data make accurate assessment of correlations challenging. This study provides a toolkit, SCSC (https://github.com/thecailab/SCSC), for the estimation of correlation coefficients in single-cell sequencing data. It comprehensively assessed four strategies (classical, non-zero, dropout-weighted, imputation) and the impact of data features in various simulated scenarios. The study found that filtering zeros significantly improves estimation accuracy, and further improvement can be achieved by considering the drop-out probability. In addition, the study also identified data features including expression level, library size, and biological variations that affect correlation estimation.

## Introduction

Single-cell sequencing provides efficiently quantification of changes in transcriptome, epigenome and other -omes separately or jointly at the individual cell level. It opens up new opportunities for screening associations of genomic features such as gene co-expression and histone modification-expression association. However, high levels of noise and sparsity in single cell sequencing data make it challenging to produce reliable assessment using typical correlation measures such as Pearson correlation coefficients. Especially, the high drop-out rate leaves zero values without identified sources, which may be from real biological signals or drop-out events. One potential solution is to treat zero values as missing values and remove them, which can reduce the effects of drop-out, natural variation, and other technical and biological noises, especially for features with low sequencing counts. However, this approach raises concerns about the potential loss of true biological information, which could affect estimation especially when the sample size is small. To improve the estimation, this study proposes a method that considers the probability of drop-out zero when estimating correlation coefficients.

An alternative approach is to impute the true gene expression data and use the imputed data for correlation analysis. We comprehensively evaluated the performance of these different strategies for correlation analysis of single-cell sequencing data in various scenarios to guide the analysis. A R package, SCSC (https://github.com/thecailab/SCSC), is provided for the efficient application of <u>s</u>ingle <u>c</u>ell <u>s</u>equencing data <u>c</u>orrelation analysis.

## Method

### Simulation

We simulated scRNA-seq datasets using SCRIP [1], which is based on a Gamma-Poisson hierarchical model. To generate correlated gene expression, we used a copula-based method which was implemented in the ‘simstudy’ package (https://github.com/kgoldfeld/simstudy) to simulate the multivariate Gamma distribution. We fit the scRNA-seq read counts from 80 acinar cells in a healthy subject from the E-MTAB-5061 dataset [2] to a Gamma distribution for the mean expression. We simulated various scenarios with different parameters, which are summarized in Table 1. These parameters include the correlation between genes (*ρ*), the number of cells, the library size (total counts for each cell), the degree of freedom of biological coefficient variance (which controls the variability of Biological Coefficient of Variation (BCV)), and the scale of BCV (which controls the overall BCV level). The drop-out rate (proportion of zeros from drop-out event) was simulated as 0.5.

**Table 1.** Parameter settings in simulations

| Parameter | Value |
| --- | --- |
| Mean expression | $\lambda = \text{Gamma}(\alpha, \beta)$ |
| Number of cells | 50, 100, 300, 600 |
| Library size | 10,000, 30,000, 50,000, 100,000 |
| BCV scale factor | 0.5, 1, 1.5, 2 |
| BCV DF | 2, 5, 10, 20 |

### Non-dropout probability estimation

The scRNA-seq data was modeled using a zero-inflated negative binomial (ZINB) mixture model,

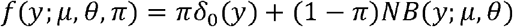

where two components were considered: Dirac delta distribution *δ*_0_ (*y*) to account for observed zeros from the drop-out, and a negative binomial distribution *NB*(*y*; *μ, θ*) models reading from actual biological material. The ZINB-WaVE [3] algorithm was employed for penalized maximum likelihood estimation of the parameters, and the posterior probability of an observation from actual biological material was calculated as 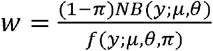.

### Correlation coefficient estimation

The sequencing reads were normalized using the size of all reads and a consistent scale factor, 10000, and transformed into the log scale. To assess the correlation of two feature vectors *x* and *y* of size *n*, three estimation strategies were formulated in the SCSC toolkit:

1. Classical estimation: The correlation coefficient was calculated using the standard Pearson formula, given by 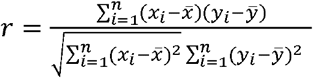.
2. Non-zero estimation: The classical estimation was performed only on the non-zero data points.
3. Dropout-weighted estimation: The weighted correlation coefficient was calculated using the 1-dropout probabilities *w*_*x*_ and *w*_*y*_, where *w*_*xi*_ and *w*_*yi*_ represent the probability that feature *x* or *y* is not a drop-out in cell *i* . The weighted correlation coefficient was given by 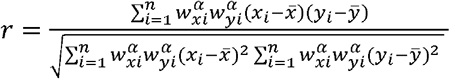. Here, the turning parameter *α* was set to 3 and the wCorr package [3] was used to perform the calculation.
4. Imputation-based estimation: Comparing to other imputation methods, SAVER [4] has an significant advantage that it removes technical variation while retaining biological variation across cells, which is critical for accurate correlation analysis [5]. Therefore, SAVER was used to impute the simulated scRNA-seq data. The uncertainty of imputed data was also considered into the correlation calculation, as proposed in [4].

The mean absolute error (MAE) were used to evaluate the performance of these methods by comparing the estimated and true correlations.

## Results

The simulation results demonstrated that filtering out zeros (non-zero estimation) significantly reduced the MAE of correlation estimation compared to using all original or imputed data, in almost all scenarios with varying drop-out rates, expression levels, total number of cells, single-cell library sizes, overdispersion scales and variations. By taking drop-out probability into consideration, the dropout-weighted estimation further improved the accuracy in most scenarios, with lower MAE (Fig 1) and standard deviation (Fig S1). Both non-zero and dropout-weighted methods produced estimation of less MAE in scenarios of higher expression level, more cells, bigger library size, or smaller biological variations.

**Figure 1.**
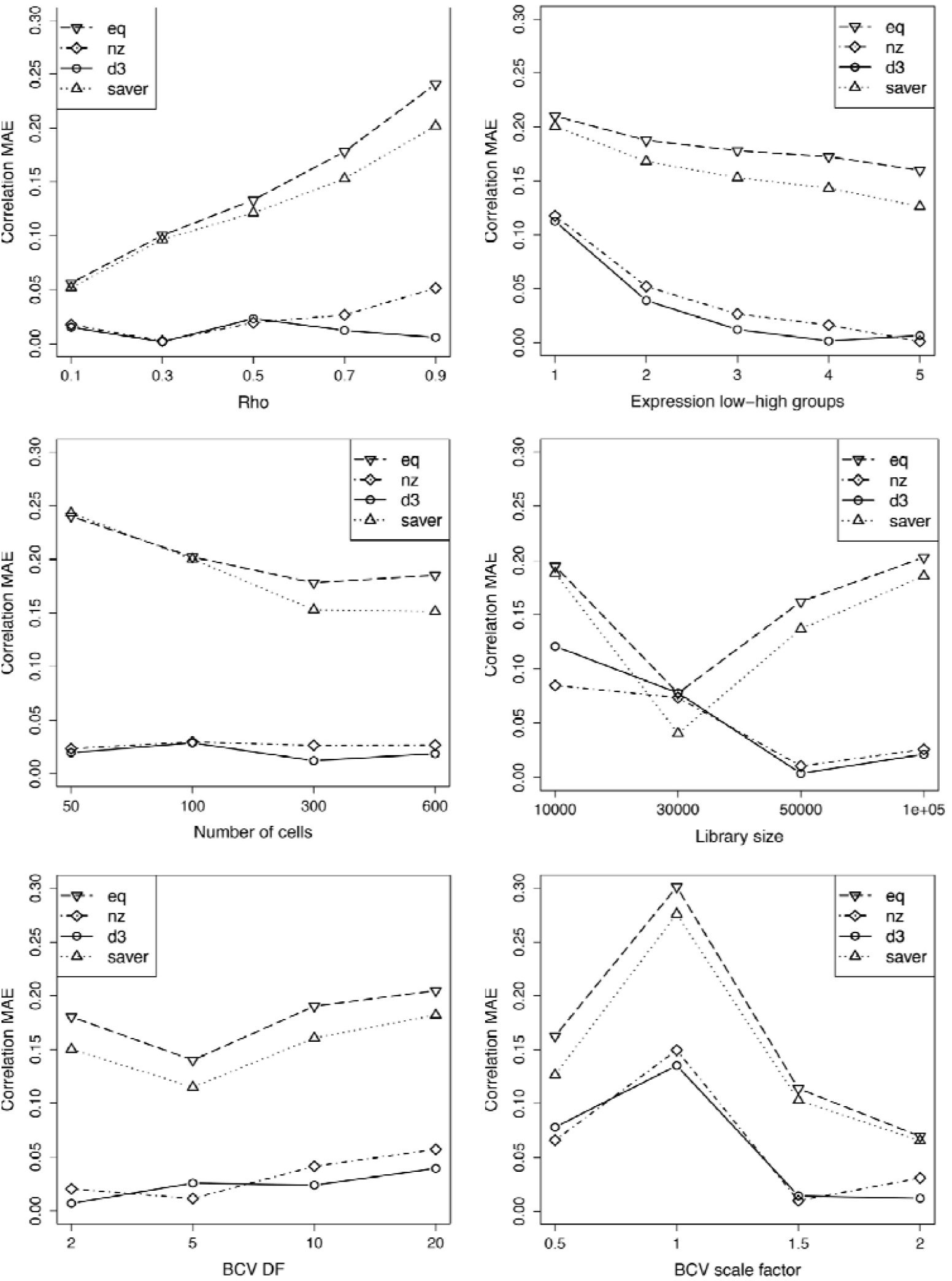
Accuracy of correlation estimation. The accuracy of correlation estimation for three methods was evaluated using the mean absolute error (MAE) metric. Simulations were performed under different scenarios with varying drop-out rates, expression level, cell numbers, library sizes, BCV level (BCV scale factor) and BCV variability (BCV DF). To compare a specific feature, other features were fixed that Rho=0.7, Expression group=3, Number of cells=300, Library size=50000, BCV df=5, and BCV scale factor=1.

In summary, this study confirmed the effectiveness of the simple zero-filtering strategy in single-cell data correlation analysis, and proposed an improved drop-out weighted approach. An R package has been developed and is avaiable for public use. The weighted method has already been successfully used to identify expression correlated super enhancers in mouse brain [6]. The SCSC package provides several major functions, including

*weight*.*calc ()* for calculating the weight factors based on drop-out probability,

*normalize*.*quant ()* for normalizing data by total counts,

*data*.*impute ()* for imputing data,

*wcorr*.*calc ()* for calculating correlation coefficients of features in two metrices, using four modes of “equal”, “nonzero”, “weighted” or “SAVER” for classical, non-zero, drop-out weighted and SAVER imputation-based estimation, respectively,

*wcorr*.*calc*.*allpairs ()* for calculating correlation coefficients of any feature pairs in one or two metrices, and

*wcorr*.*plot ()* for visualizing correlations and weights.

## Figures

**Figure S1.**
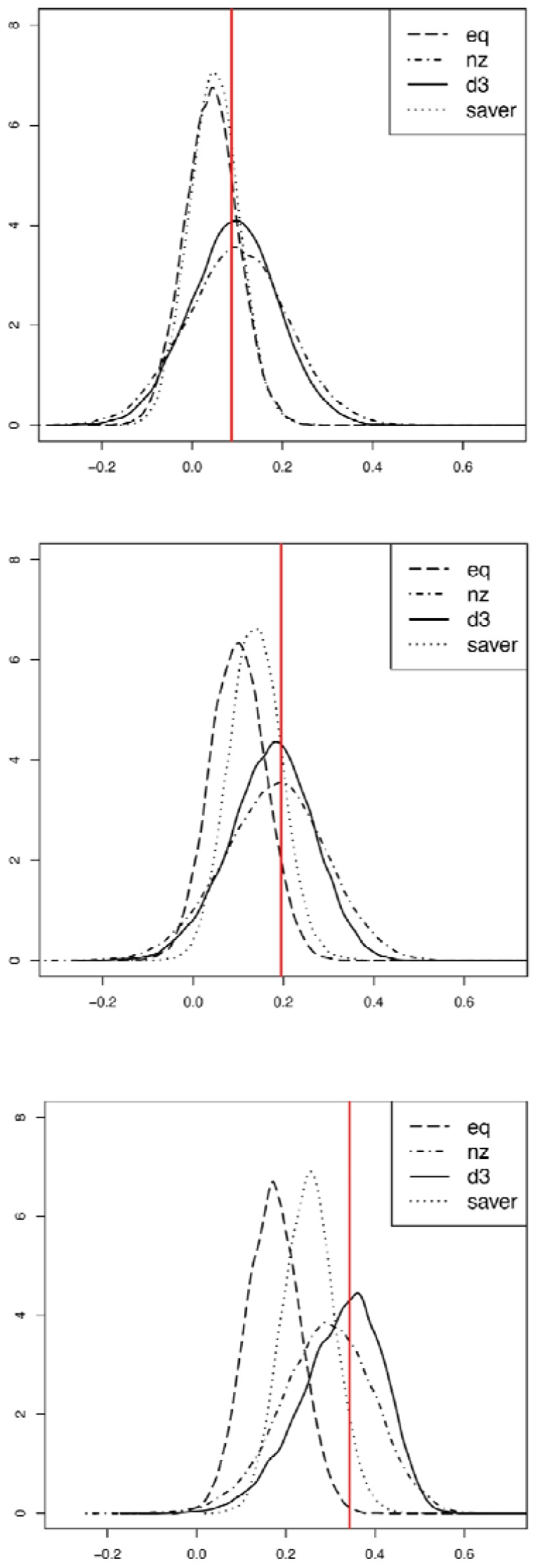
Distribution of correlation estimation. The red lines marker the mean of correlation coefficients estimated from data without technical variation.

